# Bacteria accumulate more rapidly on shorter lived flowers, but abiotic factors affect flower aging and bacterial accumulation

**DOI:** 10.1101/2025.08.19.671103

**Authors:** Rita N. Afagwu, Ciara G. Stewart, Babur S. Mirza, Avery L. Russell

## Abstract

Outcomes of ecological interactions often depend on the abundance and identity of the organisms involved. Flower-bacteria interactions can strongly affect plant ecology, and the identities of epiphytic flower bacteria are relatively well documented. Yet little is known about how the abundance of epiphytic bacteria on flowers changes over time. In this field study, we quantified how the abundance of culturable epiphytic bacteria on flowers changed as flowers aged and how abiotic factors influenced bacterial abundance and flower longevity. To accomplish this, we sampled flowers from anthesis to senescence of 8 plant species that varied substantially in terms of flower longevity and comprised 8 different genera from 7 different families. As expected, flowers of all plant species accumulated more bacteria with age. However, plant species with longer-lived flowers accumulated bacteria relatively more slowly, suggesting such plant species may have evolved more effective antibacterial defenses. Although elevated temperature is often expected to boost bacterial growth and diminish flower longevity, temperature was negatively associated with both flower longevity and bacterial accumulation, suggesting that changes to flower longevity strongly affect bacterial populations. In contrast, precipitation was positively associated with flower longevity and negatively associated with bacterial accumulation, likely because precipitation reduced plant water stress while also dislodging bacteria from flowers. Finally, we discuss the implications of our results for plant-bacterial-pollinator interactions.

## INTRODUCTION

Bacteria often shape the ecology of plants in diverse ways (Sachs et al. 2018; Compant et al. 2019; Cullen et al. 2021). For instance, beneficial soil bacteria living in association with roots can increase plant nutrient acquisition and flowering time (Lu et al. 2018; Timmusk et al. 2023) and pathogenic bacteria living in association with leaves can cause serious blights that reduce plant productivity (Guyot and Le Guen 2018). Epiphytic bacteria are particularly abundant and diverse on flowers and are thus commonly implicated in affecting flower function (Aleklett et al. 2014; Rebolleda-Gomez et al. 2019; Cullen et al. 2021). For instance, flower bacteria can cause disease, increase pollen tube growth, pollen export, and fruit set, and even produce or modify cues that shape pollinator preference (Schaeffer and Irwin 2014; Russell and Ashman 2019; Yang et al. 2019; Christensen et al. 2021; Elvira et al. 2022). However, outcomes of flower-bacteria interactions often depend on the abundance of the bacteria (Farkas et al. 2012; Junker et al. 2014; Elvira et al. 2022; Hassani et al. 2024). Because flowers are thought to accumulate bacteria as they age, the length of time the flower remains open (‘flower longevity’) might therefore strongly influence flower-bacteria interactions. In fact, flower longevity often differs substantially between plant species (< 24 hrs – 3 months; Primack 1985; Ashman and Schoen 1994; Cuartas-Domínguez et al. 2022). Yet whether differences in flower longevity affect patterns of bacterial accumulation remains poorly understood.

Bacterial populations on flowers are expected to increase with flower age for multiple reasons. Flower visitors such as butterflies, bees, and hummingbirds acquire and transfer abundant bacteria to flowers (Russell et al. 2019; Morris et al. 2020; Olson et al. 2023; Argueta-Guzman et al. 2025). Likewise, wind and rain can displace soil or flower bacteria into the air (Pietrarelli et al. 2006; Joung et al. 2017; Llontop et al. 2020) where they might be deposited on nearby flowers (Momol and Saygili 1999). Once on the flower, bacteria might also metabolize flower nutrients (e.g., petal exudates, nectar, pollen) (Vannette and Fukami 2018; Rebolleda-Gomez et al. 2019; Morales-Poole et al. 2022). Consequently, with more time for bacteria to proliferate and more chances to acquire bacteria, longer-lived flowers might ultimately acquire more bacteria. However, maintaining flowers is costly for the plant (Ashman and Schoen 1994, 1996; Cuartas-Domínguez et al. 2022), and flower bacteria often negatively affect flower function by reducing the quality of flower cues or rewards, for instance (Vannette et al. 2013; Good et al. 2014; Russell and Ashman 2019; Quevedo-Caraballo et al. 2025). We might thus expect plant taxa with longer-lived flowers to accumulate flower bacteria relatively more slowly, potentially as a consequence of greater investment in antimicrobial defenses by the plant. Alternatively, plant taxa with longer-lived flowers may have evolved to tolerate more bacteria, and so we would expect little effect of flower longevity on rates of bacterial acquisition (i.e., strategies of resistance versus tolerance, respectively; McCall and Irwin 2006; Nunez-Farfan et al. 2007). At the same time, both flower longevity and flower bacterial populations are expected to vary with abiotic conditions. For instance, higher temperatures generally reduce floral longevity by increasing metabolic activity (Hoang and Kim 2018; Arroyo et al. 2021; Song et al. 2022). In contrast, bacterial growth in flower nectar increases with increasing temperature, but growth also decreases past thermal optima (Pusey and Curry 2004; Russell and McFrederick 2022; Cecala et al. 2024). Additionally, high precipitation or humidity can delay floral senescence by reducing water stress and thus the metabolic costs of maintaining flower turgor (Hoang and Kim 2018; Dudley et al. 2018). Similarly, bacterial growth generally increases when humidity is high, as moisture availability facilitates bacterial metabolism (Pusey et al. 2009; Aleklett et al. 2014; Mahmoudi et al. 2024). As above, precipitation and wind are also expected to vector flower bacteria to flowers (Joung et al. 2017; Bhattiprolu and Monga 2018; Llontop et al. 2020), potentially increasing bacterial abundance on flowers. Yet precipitation might also dislodge epiphytic bacteria from flowers, reducing their abundance on flowers (see Butterworth and McCartney 1991). Altogether, the effects of each abiotic factor on flower bacteria abundance and flower longevity may be aligned or opposed.

In this field study, we characterized the abundance of culturable epiphytic bacteria on flowers from the onset of anthesis to senescence and how patterns of bacterial accumulation and flower longevity were influenced by abiotic factors. To accomplish this, we labeled, tracked, and collected flowers from eight plant species, representing 8 different genera and 7 different families, in a local garden across two field seasons. We hypothesized that flowers would accumulate bacteria as they aged, but that longer-lived flower types would either accumulate bacteria more slowly (potentially reflecting a strategy of resistance) or similarly (potentially reflecting a strategy of tolerance) to shorter-lived flower types. We also hypothesized that as temperature, precipitation, and humidity increased, flowers would accumulate relatively more epiphytic bacteria, given that hospitable conditions for microbes tend to increase with rising temperature and humidity, and that precipitation might vector flower bacteria. Finally, we hypothesized that as temperature increased and/or precipitation or humidity decreased, flowers would senesce more quickly, due to increased water stress.

## METHODS

### Study site and plant species selection

This field study was conducted at a community garden (Water Wise Garden, Springfield, Missouri, USA) with more than 50 flowering plant species, from July - September in 2022 and 2023. Because pollinators frequently acquire and transport floral microbes (Russell et al. 2019; Morris et al. 2020; Olson et al. 2023; Argueta-Guzman et al. 2025), we initially surveyed flowering plant species with overlapping flowering phenology (to reduce confounds with sampling different species at different times of the year) and abundant buds (to ensure sufficient flowers for sampling) for pollinator visitation. We observed abundant solitary bees, bumble bees, and butterflies on the flowers of most of these plant species. To maximize phylogenetic representation and capture a wide range of floral traits, we selected 8 co-flowering plant species, each representing a different plant genus and seven plant families (*Coreopsis verticillata*, Asteraceae; *Helianthus strumosus*, Asteraceae; *Saponaria officinalis*, Caryophyllaceae; *Commelina erecta*, Commelinaceae; *Geranium sanguineum*, Geraniaceae; *Lagerstroemia indica*, Lythraceae; *Callicarpa dichotoma*, Lamiaceae; *Eriocapitella japonica*, Ranunculaceae). In the 2022 field season, we sampled 6 plant species; to increase our sample size, in 2023 we sampled the 6 original species and an additional two species. We sampled from between 3-20 individual plants per plant species.

### Flower longevity and sampling

To track the longevity of flowers from the onset of anthesis to senescence during the study period, we used twist ties with unique identifiers to tag flower buds of each study species daily. The age of each flower was quantified in days since anthesis. Each morning at approximately the same time we inspected all study plants in the garden to identify new buds that had opened, marking anthesis. The ID of each newly opened flower bud was recorded and observed once each subsequent morning for signs of senescence. Senescence was characterized as visible signs of wilting, dropping of petals, or petal color change. We attempted to track at least n × 10 flowers for each plant species, where n = maximum flower age in days before senescing. For instance, if the flowers of a plant species opened on day 0 and the oldest flowers senesced on day 4, we attempted to track a minimum of 40 flowers in total (as flowers were open for days 0-3, with no unscenesced flowers remaining on day 4).

### Abundance of culture-dependent flower bacteria

For each plant species, each day that a flower was open, including on the day of anthesis, we collected 10-15 tagged flowers (those not exhibiting signs of senescence) to sample epiphytic bacteria. For instance, if the flowers of a given plant species senesced 4 days after anthesis, we collected a minimum of 40 tagged flowers in total (a minimum of 10 flowers of each age group; 10 flowers per day from day 0 to day 3). Flowers were deposited individually into separate sterile 15 mL centrifuge tubes (Falcon tubes, Thermo Scientific), which were stored on ice for < 2 hours before transportation to the laboratory. Strict aseptic procedures were used during the flower tagging and collection process to limit effects on flower microbial communities. In brief, we sterilized and washed scissors and forceps with 70% ethanol before and after each flower was cut and handled flowers and tags with gloves, which we periodically washed with 70% ethanol.

To detach epiphytic bacteria from floral surfaces, once flowers in centrifuge tubes were returned to the lab, 1.5 mL sterile saline (0.85% NaCl) was added and the tubes were sonicated at 40 kHz for 10 minutes (Branson M2800 Mechanical Ultrasonic Cleaner, Marshall Scientific) and then vortexed for 30 seconds. Following detachment, we plated 4 separate 100 μL aliquots from each flower sample: a “no dilution” aliquot and three serial dilution aliquots (10%, 1%, and 0.1%) on a Luria-Bertani (LB) medium often used for growing flower bacteria (e.g., Alvarez-Perez and Herrera 2013; Lee et al. 2019) with 0.02 g/ml cycloheximide to inhibit fungal growth (Vannette et al. 2012; Vannette and Fukami 2018). Plates were incubated at 30°C for 48 hours whereupon colony forming units (CFUs) were manually counted. This count was divided by the dilution factor to standardize counts across the 4 plates from each flower sample, which were then averaged and multiplied by the initial volume to estimate the total culturable bacterial population on each flower. In rare cases, plates in which most individual colonies could not be distinguished were not counted. For each sampling round we also plated a series of control plates using the sterile saline to test for contamination.

### Flower surface area

Flower surface area might influence the quantity of bacteria acquired on a flower and can influence flower longevity (Cuartas-Domínguez et al. 2022; Brazeau et al. 2024). We therefore collected 10 flowers from each of the 8 plant species in our study and flattened them with the corolla facing up, standardizing how we characterized frontal surface area of the corolla. Using an iPhone 12 camera grasped by a clamp, each flower was photographed at a standardized fixed distance, with a metric ruler in frame to provide a reference for calibration. The frontal surface area of each flower was then measured by manually tracing the corolla via the area selection tool in ImageJ (National Institutes of Health, Bethesda, MD, http://imagej.nih.gov/ij/).

### Abiotic factors

Each day of the study we recorded weather conditions (mean daily high, mean daily low, and mean daily temperature in °C, daily precipitation in cm, and mean daily dew point in °C) from the Dollison weather station (ID: KMOSPRIN363), located at 37.188° N, 93.283° W, with an elevation of 1316 feet, in Springfield, MO (Weather Underground, The Weather Company LLC). This station is situated near the field site, ensuring that the environmental data closely reflected the conditions at the study site.

### Data Analyses

All data were analyzed using R version 4.5.0 (R Development Core Team 2025). We checked assumptions of all models using the DHARMa package (Hartig 2024).

### Does floral aging influence the acquisition of epiphytic bacteria?

To assess how floral aging affected the acquisition of epiphytic floral bacteria, we used two models. To examine just the effect of flower age, we used a generalized linear mixed effects model (GLMM) with a Gaussian distribution (glmmTMB package; Magnusson et al. 2025), specifying type II Wald chi-squared ( ^2^)-tests via the Anova() function (car package; Fox and Weisberg 2019). We specified ‘total bacteria’ (estimated CFUs from each flower) as the response variable and explanatory variables as ‘plant species’ (*C. verticillata*, *H. strumosus*, *S. officinalis*, *C. erecta*, *G. sanguineum*, *L. indica*, *C. dichotoma*, or *E. japonica*) and ‘flower age’ (age of the flower in days from onset of anthesis), with ‘flower age’ within ‘plant species’ and ‘collection day’ (in days from the first of the year) within ‘field year’ (2022 or 2023) as nested random effects. To meet model assumptions, we log transformed the response variable. To examine the effect of flower longevity, we used a linear model (LM) via the lm() function and, as the response variable, specified the estimates of the coefficient regressions (‘coefficient regression estimates’) from the above GLMM of the ‘flower age’ × ‘plant species’ explanatory variable (the rate of bacterial accumulation). We specified ‘mean flower longevity’ (of each plant species) as the explanatory variable.

### Do abiotic factors influence flower epiphytic bacterial population size?

To assess how abiotic factors influenced the quantity of epiphytic bacteria found on flowers and to account for multicollinearity among multiple environmental factors, we condensed this dataset via a principal components analysis (PCA; via the prcomp() function in Base R). The PCA included ‘mean daily high temperature’, ‘mean daily low temperature’, ‘mean daily temperature’, ‘total precipitation’, and ‘mean daily dew point’ (daily mean from flower anthesis to flower collection, except for precipitation, which was a total). Before interpreting the principal component axes, we used a permutation-based test (PBT) (PCAtest package; Camargo 2024) to evaluate the overall significance of the PCA, the significance of each principal component (PC) axis, and the contributions of each variable to the significant PCs (Camargo 2022). To assess how each significant PC affected the acquisition of epiphytic floral bacteria, we next performed a principal component regression (PCR; see Hadi and Ling 1998) via the lm() function in base R. We specified ‘total bacteria’ as the response variable and each significant PC as an explanatory variable and again used type II Wald chi-squared (χ^2^)-tests to examine statistical significance. To meet model assumptions, we log transformed the response variable.

### Do abiotic factors influence flower longevity?

To assess how abiotic factors influenced floral longevity, we condensed this dataset via a PCA as above. The PCA included ‘mean daily high temperature’, ‘mean daily low temperature’, ‘mean daily temperature’, ‘total precipitation’, and ‘mean daily dew point’ (from the onset to the end of flower anthesis) and we again used a permutation-based test to interpret the statistical significance of the overall PCA and each axis and variable. We followed this by performing a PCR, specifying variables as above.

### Does flower surface area influence epiphytic bacterial population size or flower longevity?

Because flower surface area consistently differed between plant species, we assessed the role of this trait in plant species-specific patterns of epiphytic bacterial population size and flower longevity. To accomplish this, we used LMs and specified ‘initial bacteria abundance’ (mean for each plant species on the day of anthesis), ‘total bacteria abundance’ (mean for each plant species across all days), or ‘mean flower longevity’ (for each plant species in days) as the response variable and ‘mean flower surface area’ (for each plant species) as the explanatory variable. To meet model assumptions, we log transformed the bacteria abundance response variables and the explanatory variable.

## RESULTS

Flowers of all plant species accumulated significantly more culturable epiphytic bacteria as they aged (Figure 1a; GLMM: flower age effect: χ^2^_1_ = 11.367, *P* = 0.0007) and different plant species hosted significantly different quantities of epiphytic bacteria (plant species effect: χ^2^_7_ = 126.368, *P* < 0.0001). However, how quickly flowers accumulated bacteria depended significantly on plant species (Figure 1a; GLMM: flower age × plant species effect: χ^2^_7_ = 15.016, *P* = 0.0358). In fact, plant species whose flowers senesced more quickly, accumulated epiphytic floral bacteria significantly more quickly (Figure 1b; LM: F_1,6_ = 8.482, *P* = 0.0269, R^2^ = 0.517).

**Figure 1.**
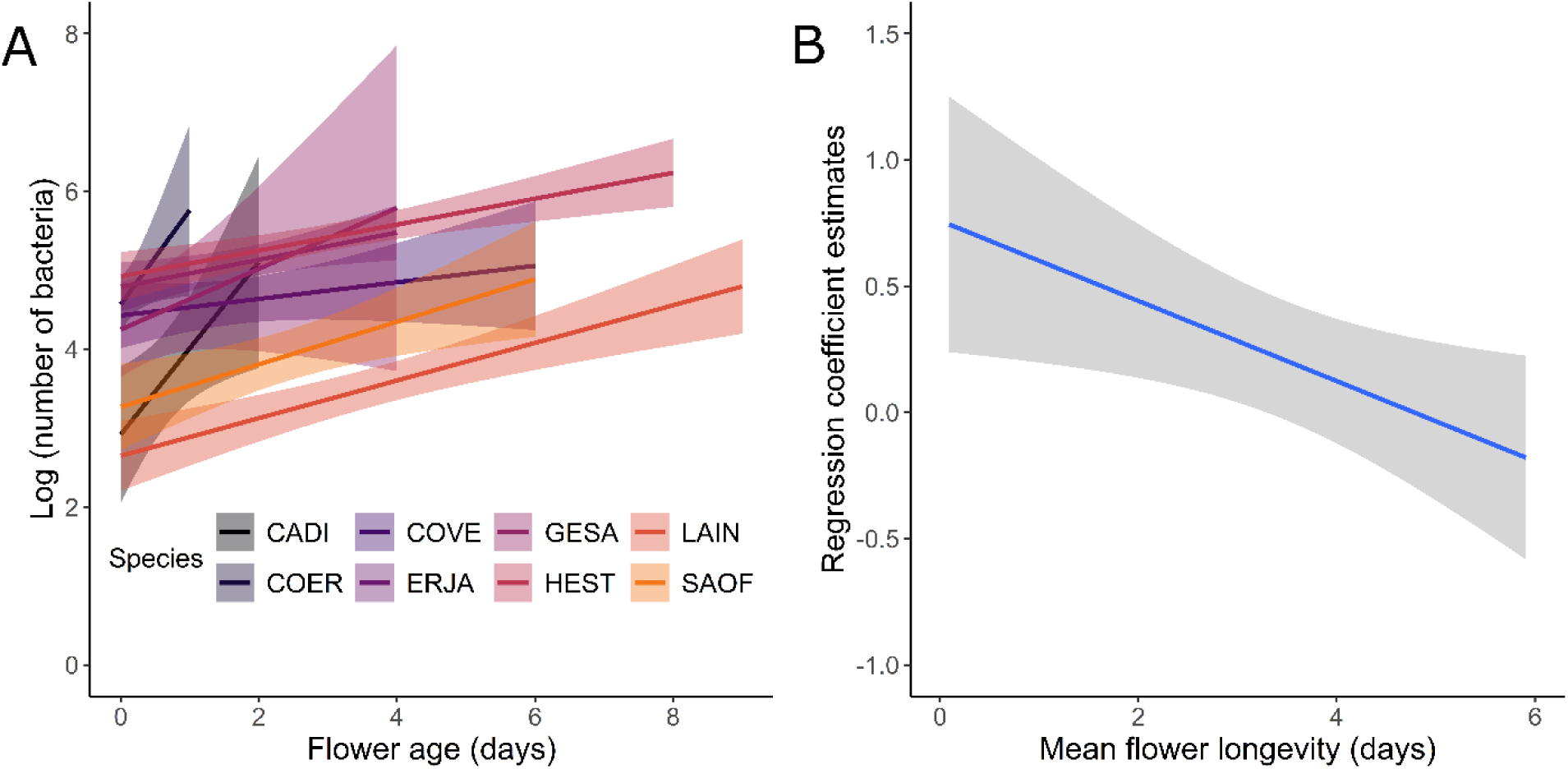
The influence of flower age and longevity, respectively, on the **A)** quantity of culturable epiphytic bacteria on flowers and **B)** rate at which those bacteria accumulate on flowers. In panel **A**, plant species are color coordinated, and abbreviations comprise the first two letters of the genus and of the species: *E. japonica* (ERJA), *H. strumosus* (HEST), *C. verticillata* (COVE), *L. indica* (LAIN), *S. officinalis* (SAOF), *G. sanguineum* GESA), *C. erecta* (COER) (Commelinaceae), and *C. dichotoma* (CADI). *N* = 360 total flowers: 52 (ERJA), 67 (HEST), 49 (COVE), 80 (LAIN), 67 (SAOF), 15 (GESA), 11 (COER), and 19 (CADI) flowers, respectively. In panel **B**, the estimates of the regression coefficients for flower age × plant species are from the model in panel **A**. Plotted lines indicate estimated means and shaded regions indicate standard errors.

The PCA investigating how abiotic factors influenced flower bacterial abundance had a non-random correlational structure, indicating it was biologically meaningful (Figure 2a: PBT: ψ = 8.476, max null = 0.132, min null = 0.011, *P* = 0; φ = 0.651, max null = 0.081, min null = 0.023, *P* = 0). This PCA yielded two statistically significant principal components (PCs) with eigenvalues > 1 that explained 92.63% of the total variation (Table 1; PBT: PC1: eigenvalue = 3.469, max null eigenvalue = 1.279, *P* = 0; PC2: eigenvalue = 1.162, max null eigenvalue = 1.178, *P* = 0.01). The first PC explained 69.39% of the variance and was significantly linked to mean daily high, low, and average temperature, as well as mean daily dew point, indicating it represents a general temperature-humidity gradient. The second PC explained 23.24% of the variance and was significantly linked to total precipitation. Bacterial abundance declined significantly as either the temperature-humidity or precipitation gradients increased (Figure 2b,c: PCR: PC1 effect: *F*_1_ = 7.380, *P* = 0.00692; PC2 effect: *F*_1_ = 79.383, *P* < 0.0001).

**Figure 2.**
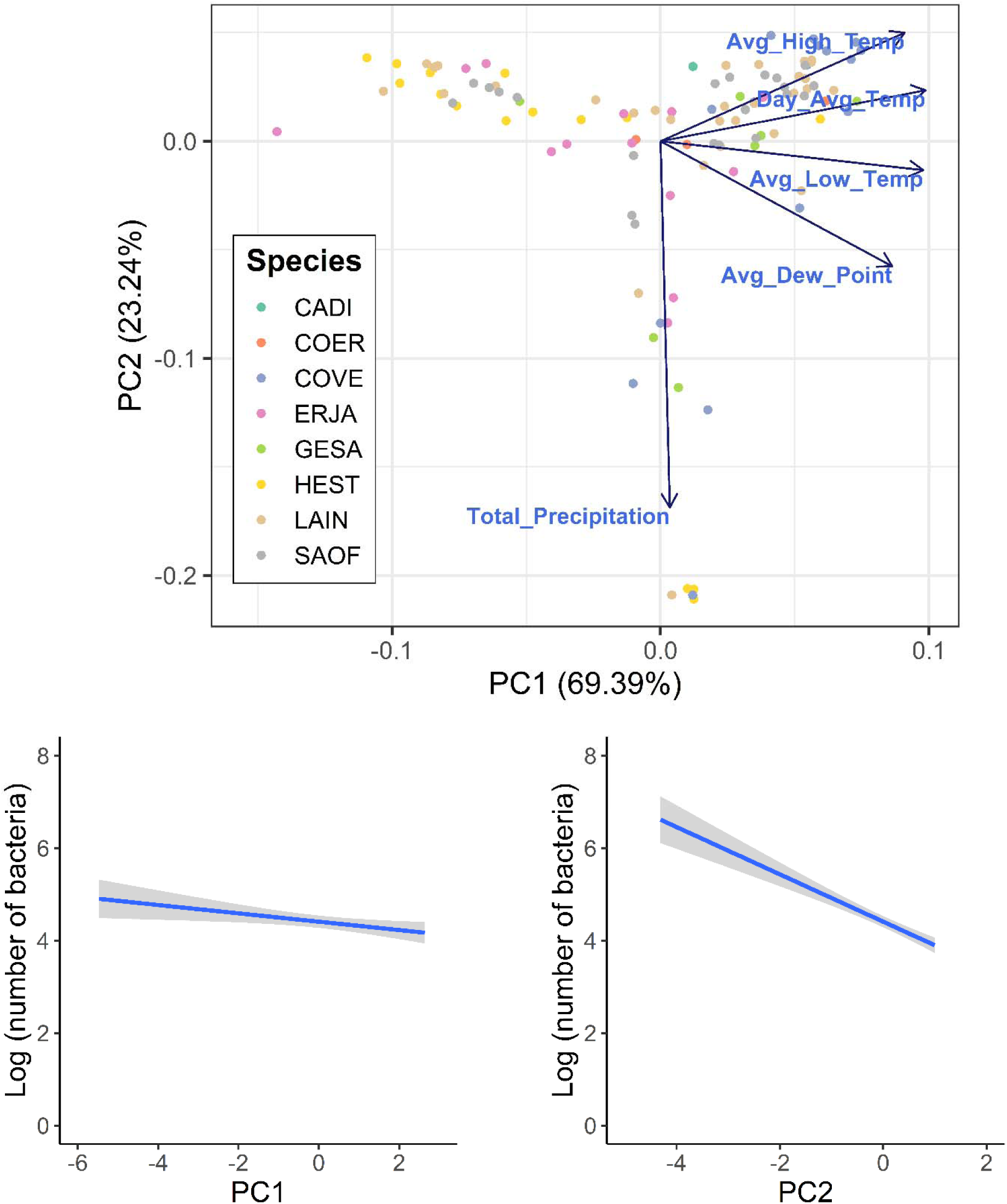
Influence of abiotic factors (temperature, humidity, and precipitation variables) on the abundance of epiphytic flower bacteria across eight plant species. **A)** Principal Component Analysis performed on five abiotic variables in which the first two principal components (PC1 and PC2) explain 69.39% and 23.24% of the variation, respectively. Arrows indicate the direction and magnitude of each environmental variable’s contribution to the principal components, with longer arrows representing stronger influences. Datapoints of the same color are the same plant species, and abbreviations comprise the first two letters of the genus and of the species: *E. japonica* (ERJA), *H. strumosus* (HEST), *C. verticillata* (COVE), *L. indica* (LAIN), *S. officinalis* (SAOF), *G. sanguineum* GESA), *C. erecta* (COER) (Commelinaceae), and *C. dichotoma* (CADI). The influence of **B)** PC1, the temperature-humidity gradient, and **C)** PC2, the precipitation gradient, on the abundance of epiphytic floral bacteria across all plant species. *N* = 360 total flowers: 52 (ERJA), 67 (HEST), 49 (COVE), 80 (LAIN), 67 (SAOF), 15 (GESA), 11 (COER), and 19 (CADI) flowers, respectively. Plotted lines indicate estimated means and shaded regions indicate standard errors.

**Table 1.** Loading of the abiotic factors on the 5 resulting principal components for the PCA investigating how abiotic factors influenced flower bacterial abundance. PCs with eigenvalues >1 indicated in bold.

|  | **PC1** | **PC2** | PC3 | PC4 | PC5 |
| --- | --- | --- | --- | --- | --- |
| Eigenvalue | **3.469** | **1.162** | 0.292 | 0.071 | 0.006 |
| Cumulative %<br>Variation | **69.386** | **23.236** | 5.836 | 1.423 | 0.119 |
| Avg_High_Temp | **-0.486** | **0.267** | -0.538 | 0.430 | 0.468 |
| Day_Avg_Temp | **-0.527** | **0.125** | -0.209 | -0.053 | -0.812 |
| Avg_Low_Temp | **-0.522** | **-0.071** | 0.125 | -0.766 | 0.346 |
| Total_Precipitation | **-0.018** | **-0.902** | -0.431 | 0.018 | -0.017 |
| Avg_Dew_Point | **-0.461** | **-0.308** | 0.682 | 0.475 | 0.045 |

The PCA investigating how abiotic factors influenced flower longevity was also biologically meaningful (Figure 3a: PBT: ψ = 7.69, max null = 0.124, min null = 0.015, *P* = 0; φ = 0.620, max null = 0.079, min null = 0.027, *P* = 0). There were two statistically significant PCs with eigenvalues > 1 that explained 91.45% of the total variation (Table 2; PBT: PC1: eigenvalue = 3.315, max null eigenvalue = 1.258, *P* = 0; PC2: eigenvalue = 1.257, max null eigenvalue = 1.132, *P* = 0). The first PC explained 66.30% of the variance and was significantly linked to mean daily high, low, and average temperature, as well as mean daily dew point, representing a general temperature-humidity gradient. The second PC explained 25.15% of the variance and was significantly linked to total precipitation. Flower longevity was significantly reduced as the temperature-humidity gradient increased, but longevity significantly increased as the precipitation gradient increased (Figure 3b,c: PCR: PC1 effect: *F*_1_ = 4.954, *P* = 0.0266; PC2 effect: *F*_1_ = 59.958, *P* < 0.0001). Mean flower surface area per plant species did not influence the mean initial or overall mean quantity of epiphytic floral bacteria (LMs: initial bacteria abundance: *F*_1,6_ = 2.669, *P* = 0.153, R^2^ = 0.193; total bacteria abundance: *F*_1,6_ = 4.013, *P* = 0.092, R^2^ = 0.301) or flower longevity (LM: *F*_1,6_ = 3.310, *P* = 0.119, R^2^ = 0.248).

**Figure 3.**
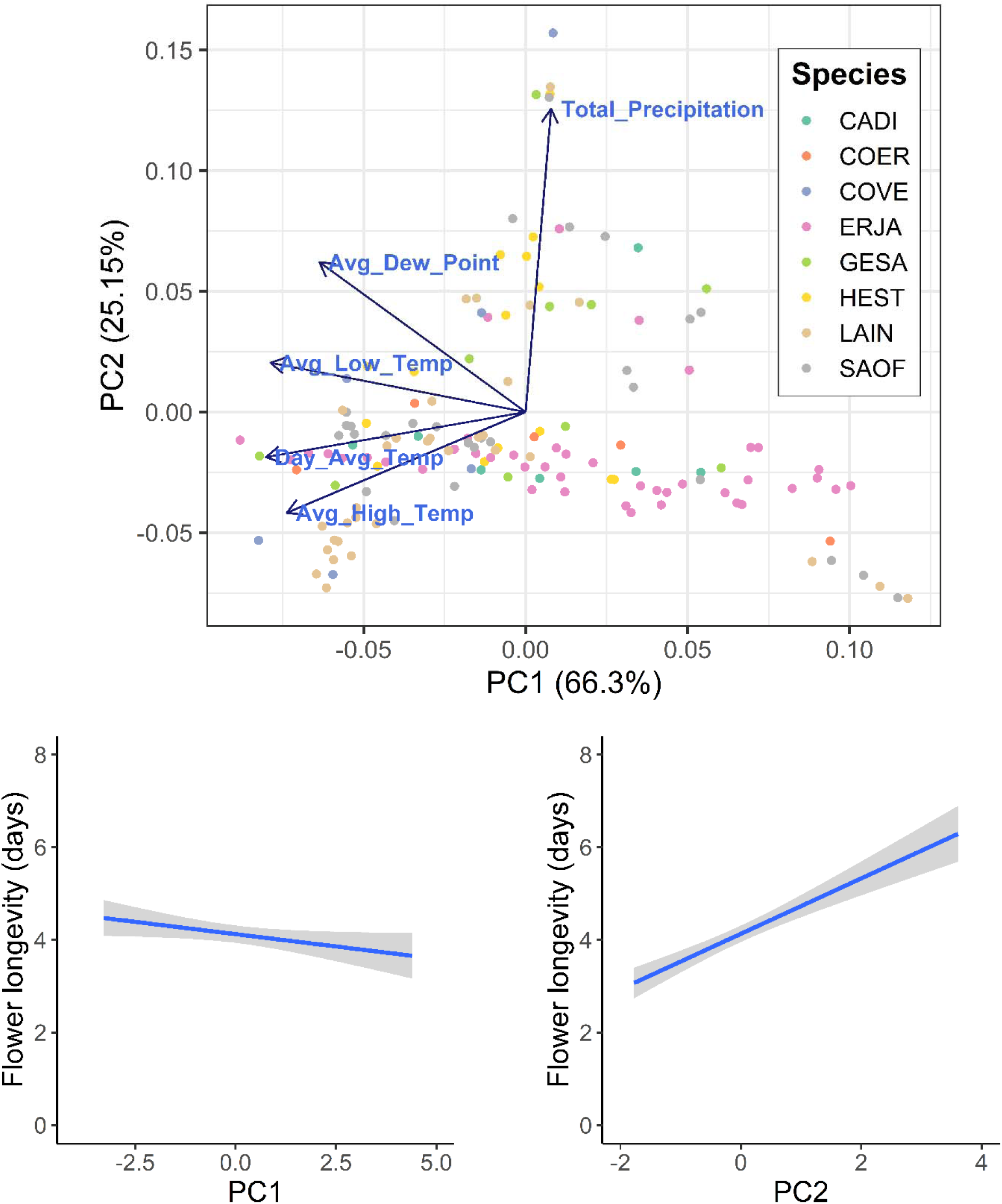
Influence of abiotic factors (temperature, humidity, and precipitation variables) on flower longevity across eight plant species. **A)** Principal Component Analysis performed on five abiotic variables in which the first two principal components (PC1 and PC2) explain 66.30% and 25.15% of the variation, respectively. Arrows indicate the direction and magnitude of each environmental variable’s contribution to the principal components, with longer arrows representing stronger influences. Datapoints of the same color are the same plant species, and abbreviations comprise the first two letters of the genus and of the species: *E. japonica* (ERJA), *H. strumosus* (HEST), *C. verticillata* (COVE), *L. indica* (LAIN), *S. officinalis* (SAOF), *G. sanguineum* GESA), *C. erecta* (COER) (Commelinaceae), and *C. dichotoma* (CADI). The influence of **B)** PC1, the temperature-humidity gradient, and **C)** PC2, the precipitation gradient, on flower longevity across all plant species. *N* = 419 total flowers: 103 (ERJA), 33 (HEST), 16 (COVE), 50 (LAIN), 156 (SAOF), 24 (GESA), 11 (COER), and 26 (CADI) flowers, respectively. Plotted lines indicate estimated means and shaded regions indicate standard errors.

**Table 2.** Loading of the abiotic factors on the 5 resulting principal components for the PCA investigating how abiotic factors influenced flower longevity. PCs with eigenvalues >1 indicated in bold.

|  | **PC1** | **PC2** | PC3 | PC4 | PC5 |
| --- | --- | --- | --- | --- | --- |
| Eigenvalue | **3.315** | **1.257** | 0.342 | 0.076 | 0.010 |
| Cumulative %<br>Variation | **66.297** | **25.147** | 6.838 | 1.511 | 0.207 |
| Avg_High_Temp | **-0.495** | **0.280** | -0.458 | 0.433 | 0.528 |
| Day_Avg_Temp | **-0.538** | **0.125** | -0.187 | 0.089 | -0.807 |
| Avg_Low_Temp | **-0.529** | **-0.137** | 0.008 | -0.802 | 0.241 |
| Total_Precipitation | **0.053** | **-0.844** | -0.524 | 0.094 | -0.034 |
| Avg_Dew_Point | **-0.427** | **-0.417** | 0.693 | 0.390 | 0.103 |

## DISCUSSION

From the initiation of anthesis to its end, on average flowers accumulated ∼10-100x as many culturable epiphytic bacteria. Although we surveyed only eight plant species, this group was taxonomically diverse, suggesting that the accumulation of epiphytic bacteria as flowers age may be a widespread pattern. At the same time, we found that floral bacteria accumulated more slowly for plant species whose flowers were longer lived. This result is consistent with a hypothesis that plant species whose flowers live relatively longer tend to evolve stronger resistance to colonization or growth of bacteria (see also McCall and Irwin 2006; Stephenson 2012). The predicted relationship between antibacterial defenses and flower longevity would be expected if larger bacterial populations on flowers generally reduce plant fitness or if negative consequences of antagonistic bacterial colonists generally increase with flower longevity (Shykoff 1996; Marre et al. 2023; Elvira et al. 2022). Additionally, we found that both flower longevity and the accumulation of flower bacteria were influenced by abiotic factors. Flower longevity across all plant species decreased as the temperature-humidity gradient increased, but floral longevity increased with greater precipitation. This result aligns well with prior findings that flower longevity is strongly negatively affected by water stress (e.g., Hoang and Kim 2018; Dudley et al. 2018). In contrast, both increased precipitation and temperature/humidity reduced the quantity of bacteria on flowers.

Aging flowers likely accumulated epiphytic bacteria because abiotic factors and flower visitors vectored bacteria onto flower surfaces (Momol and Saygili 1999; Bhattiprolu and Monga 2018; Russell et al. 2019; Argueta-Guzman et al. 2025) and because bacteria were able to use flower exudates to proliferate (Vannette and Fukami 2018; Morales-Poole et al. 2022). Given that flowers of all surveyed plant species accumulated bacteria, the rate at which bacteria die on flowers or are removed from flowers must generally be lower than the rate at which bacteria grow on or are vectored to flowers. Accordingly, older flowers – and by extension, plant species with longer-lived flowers – might contribute disproportionately to the transmission of epiphytic bacteria among flowers and flower visitors alike. Consequently, in addition to flower morphology and reward characteristics (Bodden et al. 2019; Russell et al. 2019; Adler et al. 2021), flower longevity is likely a key trait mediating bacterial transmission. Nonetheless, our results should be interpreted with some caution, given that culture-dependent methods can introduce biases by capturing only a fraction of the culturable bacteria, as found when sampling gut and soil bacteria (e.g., Fenske et al. 2020; Youseif et al. 2022). However, recent work by Hayes et al. (2021) suggests that in the case of epiphytic floral bacteria, culturing on nutrient-rich growth media recovers the most abundant flower bacterial genera (see also Morris et al. 2020), while the remaining bacterial taxa are related to rare amplicon sequence variants (ASVs). Given that our study concerns overall patterns of bacterial abundance on flowers, failure to culture rare bacteria taxa should not affect the observed patterns.

In addition to extending our understanding of flowers as hotspots of bacterial transmission (Rering et al. 2017; McFrederick and Rehan 2019; Adler et al. 2021), our study has broad implications for how floral longevity influences the function of floral displays. Assuming that the population of metabolically active bacteria increases with flower age, cues produced by flower bacteria (Russell and Ashman 2019; Yang et al. 2019; Rering et al. 2020) should also become stronger as flowers age. As a result, bacterial cues of aging flowers could more readily interfere with or complement endogenous floral cues (Russell and Ashman 2019) and might even be used by pollinators to discriminate older from younger flowers, as with age-related changes to floral color or scent (Weiss 1995; Weiss and Lamont 1997; Burdon et al. 2020). However, given that culture-dependent methods do not prevent spores from germinating, we cannot be certain that cultured bacteria were metabolically active before we removed them from the flower surface. Furthermore, no published research to our knowledge has examined what proportion of the epiphytic floral microbiome is metabolically active. Nonetheless, drawing parallels from the much more substantial phyllosphere (leaf microbiome) literature may be reasonable, given broad similarities between leaf and flower microenvironments, interactions with bacteria, and bacterial community composition (Aleklett et al. 2014; Massoni et al. 2020). Accordingly, given that metabolic activity of bacteria on leaves is common despite strong resource limitations (Moitinho et al. 2020; Thomas et al. 2024), it may be reasonable to assume that bacteria on flowers are also commonly metabolically active.

Our results also offer preliminary but compelling evidence that abiotic factors can affect floral longevity and bacterial accumulation in conflicting or complementary ways. For instance, increased precipitation was associated with greater flower longevity, which would be expected to result in greater overall accumulation of flower bacteria as above. Likewise, precipitation has been suggested to increase bacterial growth by reducing water stress, but precipitation is also thought to dislodge bacteria from flower surfaces (Momol and Saygili 1999; Llontop et al. 2020). Given that increased precipitation reduced the quantity of bacteria on flowers, the effect of dislodging epiphytic bacteria from flowers was likely far greater than potential beneficial effects on bacterial populations of greater flower longevity and reduced water stress. In contrast, the temperature-humidity gradient negatively influenced both flower longevity and the quantity of epiphytic floral bacteria. This was surprising, because increasing temperature generally increases bacterial metabolism and thus would be expected to increase bacterial abundance (Pusey and Curry 2004; Russell and McFrederick 2022; Cecala et al. 2024). One possible explanation for this negative relationship is that because higher temperatures reduced flower longevity, bacteria also had less time to accumulate on the flower surface. Additionally, heat stress inhibits photosynthesis and reduces the availability of floral metabolites and water (Marias et al. 2017; Borghi et al. 2019), presumably limiting the growth and survival of metabolically active epiphytic bacteria.

In conclusion, the community composition of flower epiphytic bacteria has been characterized by many studies (e.g., Lee et al. 2019; Hayes et al. 2021; Wei et al. 2021) and these communities are even known to change with flower age (e.g., Shade et al. 2013; Longa et al. 2022; Schaeffer et al. 2022). Our study extends our understanding of the dynamic nature of flower bacterial populations and underlines the importance of exploring how changes in flower bacterial abundance influence plant fitness, health, and plant-pollinator interactions (see also Farkas et al. 2012; Junker et al. 2014). Additionally, although we likely sampled bacteria from floral nectar, the most abundant constituents of our cultures were likely epiphytic bacteria from other parts of the flower (see also Schaeffer et al. 2022). Consistent with this, two of our eight plant species (*L. indica* and *E. japonica*) offer no nectar and yet had quantities of flower bacteria comparable to the other plant species across anthesis. Furthermore, while our study concerns flower bacteria, epiphytic flower yeast also affect plant ecology (Aleklett et al. 2014; Rebolleda-Gomez et al. 2019; Klaps et al. 2020; Deng et al. 2024). However, the relative abundance of bacteria vs yeast on flowers and how flower yeast abundance changes over time is largely unknown (but see Schaeffer et al. 2022). Finally, the evolution and plasticity of flower longevity are thought to reflect the costs of flower maintenance and variation in the pollination environment (see Xu and Servedio 2021), both of which could potentially also be affected by the accumulation of microbes.

## Supporting information

R script file zipped

## ACKNOWLEDGEMENTS

We are grateful to the Water Wise Garden naturalists for allowing us to conduct this field study and to Russell lab members and Department of Biology faculty for discussion. We acknowledge this work was performed on unceded traditional territory of the Kiikaapoi, Sioux, and Osage.

## ETHICS APPROVAL

Not applicable

## CONSENT TO PARTICIPATE

Not applicable

## CONSENT FOR PUBLICATION

Not applicable

## AVAILABILITY OF DATA AND MATERIAL

The datasets supporting this article are available as electronic supplementary material.

## CODE AVAILABILITY

The R script supporting this article is available as electronic supplementary material.

