## Supplementary material for "Bacteria accumulate more rapidly on shorter lived flowers, but abiotic factors affect flower aging and bacterial accumulation": R script file zipped: ANALYSIS_flower-age-bacteria-abund_analysis_7_1_2025.R

```
# Changelog ####
# Changed precipitation to cm

  
# Loading Packages
library(Rmisc) # Necessary for summarySE
library(readxl) # Necessary for `read_excel` function.
library(dplyr) # Necessary for group_by
library(ggplot2) # Necessary for ggplots
library(ggfortify) # Necessary for PCAs
library(kableExtra)
library(ggrepel) # Necessary for repelled labels.
library(effects) # Necessary for allEffects
library(emmeans) # Necessary for emmeans function.
library(mgcv) # Necessary for `gam` function. 
library(glmmTMB) # Necessary for `glmmTMB` function.
library(conover.test) # Necessary for conover.test
library(car) # Necessary for `vif` function.
library(lme4) # Necessary for glmer function. 
library(lmerTest)
library(glmmTMB) # Necessary for glmmTMB
library(ggpubr) # Necessary for ggarrange
library(png) # Necessary for png function
library(report) # Necessary for report function
library(DHARMa) # Model assumptions
library(PCAtest) # PCA test
library(MASS) # For PCA test

#install.packages("devtools")
#devtools::install_github("arleyc/PCAtest")

# Report
# We will use the `report` and `sessionInfo` functions to obtain all the bibliographic information we need about the R packages used in this analysis.
#report(sessionInfo())

# Name of data file: 
FILENAME1 <- "STATS_Bacteria-abundance_Flower-age_6-9-2025.csv"
FILENAME2 <- "STATS_Flower-senescence_6-9-2025.csv"
FILENAME3 <- "STATS_Summary_Bacteria-Longevity-SurfaceArea_6-1-2025.csv"

# Sets working directory automatically to wherever R script is saved
setwd(dirname(rstudioapi::getActiveDocumentContext()$path))

# Set the directory your plots will be in:
PLOTDIR <- dirname(rstudioapi::getActiveDocumentContext()$path)

setwd(dirname(rstudioapi::getActiveDocumentContext()$path)); dataset1 <- read.table(FILENAME1, sep=",", header=TRUE)
setwd(dirname(rstudioapi::getActiveDocumentContext()$path)); dataset2 <- read.table(FILENAME2, sep=",", header=TRUE)
setwd(dirname(rstudioapi::getActiveDocumentContext()$path)); dataset3 <- read.table(FILENAME3, sep=",", header=TRUE)


# Question 1: Does bacteria abundance increase with flower age? ####

## Run the model
BacAbundAge <- glmmTMB(log10(Total_Bacteria) ~ Flower_ID * Collect_Age + (1 | Flower_ID / Collect_Age ) + (1 | Field_Year / Collect_Date_Day), data = dataset1)

summary(BacAbundAge)

# Effect size calculation
emm <- emmeans(BacAbundAge, "Flower_ID")
eff_size(emm, sigma = sigma(BacAbundAge), edf = df.residual(BacAbundAge))

# Type 2 sums of squares, for Wald Test reporting.
# Note the capital Anova.
Anova(BacAbundAge)

# Response: log10(Rounded_Total_Bacteria)
#                         Chisq Df Pr(>Chisq)    
# Flower_ID             126.367  7  < 2.2e-16 ***
# Collect_Age            11.368  1  0.0007474 ***
# Flower_ID:Collect_Age  15.016  7  0.0357917 * 

# Tukey post-hoc test
summary(emmeans(BacAbundAge, pairwise ~ Flower_ID, adjust = "tukey"))


# Model assumptions via DHARMa
simulationOutput1 <- simulateResiduals(fittedModel = BacAbundAge, n = 10000)
plot(simulationOutput1) # Plot of QQ plot looks good
testUniformity(simulationOutput = simulationOutput1) # Uniformity good
testDispersion(simulationOutput1) # Dispersion good 
testZeroInflation(simulationOutput1) # Zero inflation good


## Plot the model ####

# Without predictions

#Plot model with SE
Figure1 <- ggplot(dataset1, aes(x=Collect_Age, y=log10(Total_Bacteria), color=Flower_ID, fill=Flower_ID)) + 
  geom_smooth(method = "lm", se = TRUE) + 
  ylab("Log (number of bacteria)") + xlab("Flower age (days)") + theme_classic() + 
  theme(text=element_text(size=14)) +
  scale_color_viridis_d(name = "Species", end = 0.7, option = "inferno") + 
  scale_fill_viridis_d(name = "Species", end =0.7, option = 'inferno') +
  theme(legend.position  =c(0.5, 0.2), legend.direction = "horizontal", 
        legend.background = element_rect(fill="transparent", colour = NA),
        strip.background = element_blank(),
        legend.title = element_text(size = 12), legend.text = element_text(size = 12),
        panel.spacing.x = unit(1, "lines"), 
        plot.margin = margin(c(0.2,0.51, 0.2, 0.2), unit = "cm")) + 
  scale_x_continuous(breaks = seq(0, 10, 2)) + ylim(0,8)
plot(Figure1) #To see the figure in R

# Set date
DATE <- "6-1-2025"

# Figure dimensions and name
tiff(filename=paste("Fig1_BacteriaAbundance_FlowerAge_", DATE, ".tif", sep=""), units="mm", width=125, height=125, res=1200)

# Export the figure
plot(Figure1)

# Close the plotting device
setwd(PLOTDIR); 
dev.off()


# Question 2: How does the rate of bacterial growth relate to average flower longevity ####

# Using estimates of the regression coefficients from Q1 model

## Run the model
model <- lm(Avg_Senescence_Age ~ Q1EstiCoefReg, data = dataset3)

summary(model)

# Residual standard error: 1.313 on 6 degrees of freedom
# Multiple R-squared:  0.5857,	Adjusted R-squared:  0.5166 
# F-statistic: 8.482 on 1 and 6 DF,  p-value: 0.0269

# Model assumptions via DHARMa
simulationOutput1 <- simulateResiduals(fittedModel = model, n = 10000)
plot(simulationOutput1) # Plot of QQ plot looks good ish
testUniformity(simulationOutput = simulationOutput1) # Uniformity good
testDispersion(simulationOutput1) # Dispersion good 
testZeroInflation(simulationOutput1) # Zero inflation good


#Plot model with SE
Figure1b <- ggplot(dataset3, aes(x=Avg_Senescence_Age, y=Q1EstiCoefReg)) + 
  ylab("Regression coefficient estimates") + xlab("Mean flower longevity (days)") + theme_classic() + 
  geom_smooth(method = "lm", se = TRUE) + 
  theme(text=element_text(size=14)) +
  scale_color_viridis_d(name = "Species", end = 0.7, option = "inferno") + 
  scale_fill_viridis_d(name = "Species", end =0.7, option = 'inferno') +
  theme(legend.position  =c(0.5, 0.2), legend.direction = "horizontal", 
        legend.background = element_rect(fill="transparent", colour = NA),
        strip.background = element_blank(),
        legend.title = element_text(size = 12), legend.text = element_text(size = 12),
        panel.spacing.x = unit(1, "lines"), 
        plot.margin = margin(c(0.2,0.51, 0.2, 0.2), unit = "cm")) + 
  xlim(0,6) + ylim(-1,1.5)
plot(Figure1b) #To see the figure in R

# Set date
DATE <- "6-29-2025"

# Figure dimensions and name
tiff(filename=paste("Fig1b_RegCoefficients_FlowerLong_", DATE, ".tif", sep=""), units="mm", width=125, height=125, res=1200)

# Export the figure
plot(Figure1b)

# Close the plotting device
setwd(PLOTDIR); 
dev.off()


# Question 3: How is bacteria abundance influenced by weather conditions? ####

## Principal Components Analysis
dataset1_numeric <- dataset1 %>%
  dplyr::select(Avg_High_Temp, Day_Avg_Temp, Avg_Low_Temp, Total_Precipitation, Avg_Dew_Point) %>%
  na.omit()  # Here we use na.omit to remove rows with missing values

pca_result_V2 <- prcomp(dataset1_numeric, center = TRUE, scale. = TRUE)  # Here we use prcomp to perform PCA with 

dataset1_scaled <- dataset1 %>%
  dplyr::select(Flower_ID, Avg_High_Temp, Day_Avg_Temp, Avg_Low_Temp, Total_Precipitation, Avg_Dew_Point) %>%
  na.omit()  # Here we use na.omit to match rows with pca_result_V2 for accurate coloring


# We will call this Version 2, as it is using slightly different data.
Figure2a <- autoplot(pca_result_V2, data = dataset1_scaled, 
                        colour = 'Flower_ID',  # Here we use Flower_ID for point coloring
                        loadings = TRUE, loadings.label = TRUE, loadings.color = "midnightblue", 
                        loadings.label.hjust = 1,
                        loadings.label.vjust = 1,
                        loadings.label.colour = "royalblue",
                        loadings.label.fontface = "bold") +
  scale_color_brewer(palette = "Set2", name = "Species") + 
  theme_bw(base_size = 15) +
  theme(legend.position = c(0.125,0.35),
        legend.background = element_rect(color = "black", size = 0.3),
        legend.margin = margin(5, 5, 5, 5),
        legend.title = element_text(face = "bold", hjust = 0.5),          
        legend.box.margin=margin(0, 0, 0, 0),
        legend.spacing.x = unit(0.5, 'cm')) + 
  guides(fill = guide_legend(title = "Species", title.position = "top", nrow = 8, byrow = TRUE))
Figure2a

# Set date
DATE <- "6-1-2025"

# Figure dimensions and name
tiff(filename=paste("Fig2a_PCA_Weather_BactAbund_", DATE, ".tif", sep=""), units="mm", width=150, height=125, res=1200)

# Export the figure
plot(Figure2a)

# Close the plotting device
setwd(PLOTDIR); 
dev.off()

## Eigenvalues ####

# Let's proceed by creating the Eigenvalue table that correspond this PCA.
EigenTable2A <- as.data.frame(eigen(cor(na.omit(dataset1_scaled[,-1])))$values)
EigenTable2A <- as.data.frame(EigenTable2A)
colnames(EigenTable2A) <- "Eigenvalue"

EigenTable2A$CumulativePercentVariation <- EigenTable2A$Eigenvalue/sum(EigenTable2A$Eigenvalue)*100
EigenTable2A <- t(EigenTable2A) # "t" function transposes. 

EigenTable2B <- as.data.frame(eigen(cor(na.omit(dataset1_scaled[,-1])))$vectors)
EigenTable <- rbind(EigenTable2A,EigenTable2B)

row.names(EigenTable) <-c("Eigenvalue", "Cumulative % Variation", colnames(na.omit(dataset1_scaled[,-1])))
colnames(EigenTable) <- c("PC1", "PC2", "PC3", "PC4", "PC5")

kbl(round(EigenTable,3)) %>%
  kable_material_dark()


## Interpretation ####

# Using PCAtest for statistical significance of PCs
result<-PCAtest(dataset1_numeric, 100, 100, 0.05, varcorr=FALSE, counter=FALSE, plot=TRUE)

# Test of PCA significance: 5 variables, 360 observations
# 100 bootstrap replicates, 100 random permutations

# Empirical Psi = 8.4762, Max null Psi = 0.1323, Min null Psi = 0.0105, p-value = 0
# Empirical Phi = 0.6510, Max null Phi = 0.0813, Min null Phi = 0.0229, p-value = 0

# Empirical eigenvalue #1 = 3.46931, Max null eigenvalue = 1.27866, p-value = 0
# Empirical eigenvalue #2 = 1.1618, Max null eigenvalue = 1.17783, p-value = 0.01
# Empirical eigenvalue #3 = 0.2918, Max null eigenvalue = 1.05195, p-value = 1
# Empirical eigenvalue #4 = 0.07113, Max null eigenvalue = 0.98399, p-value = 1
# Empirical eigenvalue #5 = 0.00595, Max null eigenvalue = 0.93295, p-value = 1

# PC 1 is significant and accounts for 69.4% (95%-CI:68.3-70.6) of the total variation
# PC 2 is significant and accounts for 23.2% (95%-CI:22.4-24.3) of the total variation

# 2 PC axes are significant and account for 92.6% of the total variation

# Variables 1, 2, 3, and 5 have significant loadings on PC 1
# Variable 4 has a significant loading on PC 2


#  **PC1** accounts for approximately 69.4% of the variance, primarily representing 
# temperature-related variables such as **High_Temp**, **Day_Avg_Temp**,  **Low_Temp** 
# and **Avg_Dew_Point** (with strong negative loadings), which suggests it captures a general 
# "temperature gradient." **PC2**, explaining 23.2% of the variance, is dominated by 
# **Precipitation** (strong negative loading), indicating it 
# represents a "humidity-related" component. Together, **PC1** and **PC2** explain over 
# 93% of the variation, which provides a substantial reduction in dimensionality while 
# retaining most of the information in the dataset. Subsequent components contribute 
# minimally, each accounting for less than 6% of the total variance, suggesting they 
# contain only minor patterns or noise. 


## Principal Components Regression ####
# To assess directional relationship between each PC and the response variable

# First extract the PCs (there are only 5 for this model)
pca_result_V2$x [,1:5] %>% head(1) 

# Put the PCs into a data frame
pcs <- as.data.frame(pca_result_V2$x)

# Rename the rows so that column 1 is renamed so we can merge datasets
pcs <- cbind(rownames(pcs), pcs)
rownames(pcs) <- NULL
colnames(pcs) <- c("Row1","PC1","PC2","PC3","PC4","PC5")

# Put the microbe abundance into a data frame
response <- as.data.frame(dataset1$Total_Bacteria)

# Rename the rows so that column 1 is renamed so we can merge datasets
response <- cbind(rownames(response), response)
rownames(response) <- NULL
colnames(response) <- c("Row1","Total_Bacteria")

# Combine the data frames horizontally (6 total columns = 5+1 columns)
pcs.abund <- merge(response,pcs,by="Row1")

# Plot it. See there is non-normality, so violates assumptions of LM
plot(pcs$PC1,log10(dataset1$Total_Bacteria))

# Plot it. See there is non-normality, so violates assumptions of LM
plot(pcs$PC2,log10(dataset1$Total_Bacteria))


# Linear model for PC1 and PC2
PCR.BactAbund <- lm(log10(Total_Bacteria) ~ PC1 + PC2, data = pcs.abund)

summary(PCR.BactAbund)

Anova(PCR.BactAbund)

# Response: log10(Total_Bacteria)
# Sum Sq  Df F value    Pr(>F)    
# PC1        10.14   1  7.3795  0.006918 ** 
# PC2       109.12   1 79.3833 < 2.2e-16 ***
# Residuals 490.73 357

# Model assumptions via DHARMa
simulationOutput1 <- simulateResiduals(fittedModel = PCR.BactAbund, n = 10000)
plot(simulationOutput1) # Plot of QQ plot looks good
testUniformity(simulationOutput = simulationOutput1) # Uniformity good
testDispersion(simulationOutput1) # Dispersion good 
testZeroInflation(simulationOutput1) # Zero inflation good


#Plot model with SE
Figure2b <- ggplot(pcs.abund, aes(x=PC1, y=log10(Total_Bacteria))) + 
  geom_smooth(method = "lm", se = TRUE) + 
  ylab("Log (number of bacteria)") + xlab("PC1") + theme_classic() + 
  theme(text=element_text(size=14)) +
  scale_color_viridis_d(name = "Species", end = 0.7, option = "inferno") + 
  scale_fill_viridis_d(name = "Species", end =0.7, option = 'inferno') +
  theme(legend.position  =c(0.5, 0.2), legend.direction = "horizontal", 
        legend.background = element_rect(fill="transparent", colour = NA),
        strip.background = element_blank(),
        legend.title = element_text(size = 12), legend.text = element_text(size = 12),
        panel.spacing.x = unit(1, "lines"), 
        plot.margin = margin(c(0.2,0.51, 0.2, 0.2), unit = "cm")) + 
  xlim(-6,3) + ylim(0,8)
plot(Figure2b) #To see the figure in R

# Set date
DATE <- "6-12-2025"

# Figure dimensions and name
tiff(filename=paste("Fig2b_Bacteria_PCR_PC1_", DATE, ".tif", sep=""), units="mm", width=85, height=85, res=1200)

# Export the figure
plot(Figure2b)

# Close the plotting device
setwd(PLOTDIR); 
dev.off()


#Plot model with SE
Figure2c <- ggplot(pcs.abund, aes(x=PC2, y=log10(Total_Bacteria))) + 
  geom_smooth(method = "lm", se = TRUE) + 
  ylab("Log (number of bacteria)") + xlab("PC2") + theme_classic() + 
  theme(text=element_text(size=14)) +
  scale_color_viridis_d(name = "Species", end = 0.7, option = "inferno") + 
  scale_fill_viridis_d(name = "Species", end =0.7, option = 'inferno') +
  theme(legend.position  =c(0.5, 0.2), legend.direction = "horizontal", 
        legend.background = element_rect(fill="transparent", colour = NA),
        strip.background = element_blank(),
        legend.title = element_text(size = 12), legend.text = element_text(size = 12),
        panel.spacing.x = unit(1, "lines"), 
        plot.margin = margin(c(0.2,0.51, 0.2, 0.2), unit = "cm")) + 
  xlim(-5,2) + ylim(0,8)
plot(Figure2c) #To see the figure in R

# Set date
DATE <- "6-12-2025"

# Figure dimensions and name
tiff(filename=paste("Fig2c_Bacteria_PCR_PC2_", DATE, ".tif", sep=""), units="mm", width=85, height=85, res=1200)

# Export the figure
plot(Figure2c)

# Close the plotting device
setwd(PLOTDIR); 
dev.off()


# Question 4: How is flower longevity influenced by weather conditions? ####

## PCA
# Next we perform the PCA using the prcomp function.
dataset2_numeric <- dataset2 %>%
  dplyr::select(Avg_High_Temp, Day_Avg_Temp, Avg_Low_Temp, Total_Precipitation, Avg_Dew_Point) %>%
  na.omit()  

pca_result <- prcomp(dataset2_numeric, center = TRUE, scale. = TRUE)

dataset2_scaled <- dataset2 %>%
  dplyr::select(Flower_ID, Avg_High_Temp, Day_Avg_Temp, Avg_Low_Temp, Total_Precipitation, Avg_Dew_Point) %>%
  na.omit()  

Figure3a <- autoplot(pca_result, data = dataset2_scaled, 
                     colour = 'Flower_ID',  # Use Flower_ID for coloring
                     loadings = TRUE, loadings.label = TRUE, loadings.color = "midnightblue", 
                     loadings.label.hjust = -0.05,
                     loadings.label.colour = "royalblue",
                     loadings.label.fontface = "bold") +
  scale_color_brewer(palette = "Set2", name = "Species") + 
  theme_bw(base_size = 15) +
  theme(legend.position = c(0.875,0.65),
        legend.background = element_rect(color = "black", size = 0.3),
        legend.margin = margin(5, 5, 5, 5),
        legend.title = element_text(face = "bold", hjust = 0.5),          
        legend.box.margin=margin(0, 0, 0, 0),
        legend.spacing.x = unit(0.5, 'cm')) + 
  guides(fill = guide_legend(title = "Species", title.position = "top", nrow = 8, byrow = TRUE))
Figure3a

# Set date
DATE <- "6-12-2025"

# Figure dimensions and name
tiff(filename=paste("Fig3a_PCA_Weather_FlowerSenesc_", DATE, ".tif", sep=""), units="mm", width=150, height=125, res=1200)

# Export the figure
plot(Figure3a)

# Close the plotting device
setwd(PLOTDIR); 
dev.off()


## Eigenvalues ####
# In the context of PCA, eigenvectors represent the directions (or axes) along which the data varies the most, known as the principal components. Eigenvalues, on the other hand, indicate how much of the data’s variance is captured by each of these components. The first principal component, with the largest eigenvalue, captures the most variation in the data, while each subsequent component captures less. Essentially, eigenvalues tell us how important each principal component is, and eigenvectors show the directions of the greatest variation in the data.

# Let's proceed by creating the Eigenvalue table that correspond this PCA.
EigenTable2A <- as.data.frame(eigen(cor(na.omit(dataset2_scaled[,-1])))$values)
EigenTable2A <- as.data.frame(EigenTable2A)
colnames(EigenTable2A) <- "Eigenvalue"

EigenTable2A$CumulativePercentVariation <- EigenTable2A$Eigenvalue/sum(EigenTable2A$Eigenvalue)*100
EigenTable2A <- t(EigenTable2A) # "t" function transposes. 

EigenTable2B <- as.data.frame(eigen(cor(na.omit(dataset2_scaled[,-1])))$vectors)
EigenTable <- rbind(EigenTable2A,EigenTable2B)

row.names(EigenTable) <-c("Eigenvalue", "Cumulative % Variation", colnames(na.omit(dataset2_scaled[,-1])))
colnames(EigenTable) <- c("PC1", "PC2", "PC3", "PC4", "PC5")

kbl(round(EigenTable,3)) %>%
  kable_material_dark()

## Interpretation ####

# Using PCAtest for statistical significance of PCs
result<-PCAtest(dataset2_numeric, 100, 100, 0.05, varcorr=FALSE, counter=FALSE, plot=TRUE)

# Test of PCA significance: 5 variables, 419 observations
# 100 bootstrap replicates, 100 random permutations

  
# Empirical Psi = 7.6920, Max null Psi = 0.1239, Min null Psi = 0.0150, p-value = 0
# Empirical Phi = 0.6202, Max null Phi = 0.0787, Min null Phi = 0.0273, p-value = 0

# Empirical eigenvalue #1 = 3.31486, Max null eigenvalue = 1.25837, p-value = 0
# Empirical eigenvalue #2 = 1.25737, Max null eigenvalue = 1.13192, p-value = 0
# Empirical eigenvalue #3 = 0.3419, Max null eigenvalue = 1.05185, p-value = 1
# Empirical eigenvalue #4 = 0.07554, Max null eigenvalue = 0.98917, p-value = 1
# Empirical eigenvalue #5 = 0.01034, Max null eigenvalue = 0.92974, p-value = 1

# PC 1 is significant and accounts for 66.3% (95%-CI:65-67.8) of the total variation
# PC 2 is significant and accounts for 25.1% (95%-CI:24.3-26) of the total variation

# 2 PC axes are significant and account for 91.4% of the total variation

# Variables 1, 2, 3, and 5 have significant loadings on PC 1
# Variable 4 has a significant loading on PC 2

# Interpretation: The PCA results indicate that the first principal component (PC1) 
# explains the majority of the variance (66.3%), with subsequent components explaining 
# diminishing amounts of variation: PC2 at 25.1%, PC3 at 6.48%, and so forth. PC1 is 
# primarily associated with negative loadings on High_Temp, Day_Avg_Temp, Low_Temp, 
# and Avg_Dew_Point, suggesting that this component captures an overall temperature-
# related trend. PC2, which explains 25.1% of the variance, has a strong negative loading 
# on Precipitation and a bit here of Avg_Dew_Point too, indicating that this component may 
# represent a variation driven by precipitation and some humidity.


# Suggestions for the model:

# Drop Redundant Temperature Variables: We could retain only one or two representative 
# temperature variables (e.g., Day_Avg_Temp and Low_Temp ) to capture overall temperature 
# trends, reducing multicollinearity. Dropping some of these correlated temperature 
# variables will help decrease redundancy.

# Keep Precipitation: Since Precipitation is positioned orthogonally, it provides unique 
# information and is less likely to introduce multicollinearity. Keeping it in the model 
# is beneficial for capturing precipitation-related variance.

# Since Avg_Dew_Point is another measure of atmospheric moisture that tends to correlate 
# with temperature, we could consider dropping it to reduce multicollinearity.


## Principal Components Regression ####
# To assess directional relationship between each PC and the response variable

# First extract the PCs (there are only 5 for this model)
pca_result$x [,1:5] %>% head(1) 

# Put the PCs into a data frame
pcs.2 <- as.data.frame(pca_result$x)

# Rename the rows so that column 1 is renamed so we can merge datasets
pcs.2 <- cbind(rownames(pcs.2), pcs.2)
rownames(pcs.2) <- NULL
colnames(pcs.2) <- c("Row1","PC1","PC2","PC3","PC4","PC5")

# Put the microbe abundance into a data frame
response.2 <- as.data.frame(dataset2$Senesce_Age)

# Rename the rows so that column 1 is renamed so we can merge datasets
response.2 <- cbind(rownames(response.2), response.2)
rownames(response.2) <- NULL
colnames(response.2) <- c("Row1","Senesce_Age")

# Combine the data frames horizontally (6 total columns = 5+1 columns)
pcs.senesc <- merge(response.2,pcs.2,by="Row1")

# Plot it. See there is non-normality, so violates assumptions of LM
plot(pcs.2$PC1,(dataset2$Senesce_Age))

# Plot it. See there is non-normality, so violates assumptions of LM
plot(pcs.2$PC2,(dataset2$Senesce_Age))


# Linear model for PC1 and PC2
PCR.Senesc <- lm((Senesce_Age+1) ~ PC1 + PC2, data = pcs.senesc)

summary(PCR.Senesc)

Anova(PCR.Senesc)

# Response: (Senesce_Age + 1)
# Sum Sq  Df F value    Pr(>F)    
# PC1         18.29   1  4.9542   0.02656 *  
# PC2        221.39   1 59.9577 7.452e-14 ***

# Model assumptions via DHARMa
simulationOutput1 <- simulateResiduals(fittedModel = PCR.Senesc, n = 10000)
plot(simulationOutput1) # Plot of QQ plot looks good
testUniformity(simulationOutput = simulationOutput1) # Uniformity good
testDispersion(simulationOutput1) # Dispersion good 
testZeroInflation(simulationOutput1) # Zero inflation good


#Plot model with SE
Figure3b <- ggplot(pcs.senesc, aes(x=PC1, y=Senesce_Age)) + 
  geom_smooth(method = "lm", se = TRUE) + 
  ylab("Flower longevity (days)") + xlab("PC1") + theme_classic() + 
  theme(text=element_text(size=14)) +
  scale_color_viridis_d(name = "Species", end = 0.7, option = "inferno") + 
  scale_fill_viridis_d(name = "Species", end =0.7, option = 'inferno') +
  theme(legend.position  =c(0.5, 0.2), legend.direction = "horizontal", 
        legend.background = element_rect(fill="transparent", colour = NA),
        strip.background = element_blank(),
        legend.title = element_text(size = 12), legend.text = element_text(size = 12),
        panel.spacing.x = unit(1, "lines"), 
        plot.margin = margin(c(0.2,0.51, 0.2, 0.2), unit = "cm")) + 
  xlim(-4,5) + ylim(0,8)
plot(Figure3b) #To see the figure in R

# Set date
DATE <- "6-29-2025"

# Figure dimensions and name
tiff(filename=paste("Fig3b_Senesce_PCR_PC1_", DATE, ".tif", sep=""), units="mm", width=85, height=85, res=1200)

# Export the figure
plot(Figure3b)

# Close the plotting device
setwd(PLOTDIR); 
dev.off()


#Plot model with SE
Figure3c <- ggplot(pcs.senesc, aes(x=PC2, y=Senesce_Age)) + 
  geom_smooth(method = "lm", se = TRUE) + 
  ylab("Flower longevity (days)") + xlab("PC2") + theme_classic() + 
  theme(text=element_text(size=14)) +
  scale_color_viridis_d(name = "Species", end = 0.7, option = "inferno") + 
  scale_fill_viridis_d(name = "Species", end =0.7, option = 'inferno') +
  theme(legend.position  =c(0.5, 0.2), legend.direction = "horizontal", 
        legend.background = element_rect(fill="transparent", colour = NA),
        strip.background = element_blank(),
        legend.title = element_text(size = 12), legend.text = element_text(size = 12),
        panel.spacing.x = unit(1, "lines"), 
        plot.margin = margin(c(0.2,0.51, 0.2, 0.2), unit = "cm")) + 
  xlim(-2,4) + ylim(0,8)
plot(Figure3c) #To see the figure in R

# Set date
DATE <- "6-29-2025"

# Figure dimensions and name
tiff(filename=paste("Fig3c_Senesce_PCR_PC2_", DATE, ".tif", sep=""), units="mm", width=85, height=85, res=1200)

# Export the figure
plot(Figure3c)

# Close the plotting device
setwd(PLOTDIR); 
dev.off()


# Question 5: Does surface area affect how many bacteria are on flowers or senescence  ####


# Initial quantity of bacteria
# Run the model
Day0BacAbund.SurfArea <- lm(log10(Day0_Avg_Bacteria) ~ log10(Avg_Surface_Area), data = dataset3)

summary(Day0BacAbund.SurfArea)

# Residual standard error: 0.4184 on 6 degrees of freedom
# Multiple R-squared:  0.3079,	Adjusted R-squared:  0.1925 
# F-statistic: 2.669 on 1 and 6 DF,  p-value: 0.1534

# Model assumptions via DHARMa
simulationOutput1 <- simulateResiduals(fittedModel = Day0BacAbund.SurfArea, n = 10000)
plot(simulationOutput1) # Plot of QQ plot looks good
testUniformity(simulationOutput = simulationOutput1) # Uniformity good
testDispersion(simulationOutput1) # Dispersion good 
testZeroInflation(simulationOutput1) # Zero inflation good


# Total average quantity of bacteria
# Run the model
TotBacAbund.SurfArea <- lm(log10(Total_Avg_Bacteria) ~ log10(Avg_Surface_Area), data = dataset3)

summary(TotBacAbund.SurfArea)

# Residual standard error: 0.02083 on 6 degrees of freedom
# Multiple R-squared:  0.3555,	Adjusted R-squared:  0.2481 
# F-statistic:  3.31 on 1 and 6 DF,  p-value: 0.1187

# Model assumptions via DHARMa
simulationOutput1 <- simulateResiduals(fittedModel = TotBacAbund.SurfArea, n = 10000)
plot(simulationOutput1) # Plot of QQ plot looks good
testUniformity(simulationOutput = simulationOutput1) # Uniformity good
testDispersion(simulationOutput1) # Dispersion good 
testZeroInflation(simulationOutput1) # Zero inflation good


# Senescence age vs surface area
# Run the model
Senescence.SurfArea <- lm(Avg_Senescence_Age ~ log10(Avg_Surface_Area), data = dataset3)

summary(Senescence.SurfArea)

# Residual standard error: 1.579 on 6 degrees of freedom
# Multiple R-squared:  0.4008,	Adjusted R-squared:  0.3009 
# F-statistic: 4.013 on 1 and 6 DF,  p-value: 0.092

# Model assumptions via DHARMa
simulationOutput1 <- simulateResiduals(fittedModel = Senescence.SurfArea, n = 10000)
plot(simulationOutput1) # Plot of QQ plot looks good
testUniformity(simulationOutput = simulationOutput1) # Uniformity good
testDispersion(simulationOutput1) # Dispersion good 
testZeroInflation(simulationOutput1) # Zero inflation good
```
